# The first spectrum of spore form and function reveals constrained evolution in mycorrhizal symbiosis

**DOI:** 10.64898/2026.08.21.746157

**Authors:** Carlos A. Aguilar-Trigueros, Smriti Pehim Limbu, Liam Nokes, Joanna Bergmann, Matthias C. Rillig, V. Bala Chaudhary

## Abstract

Arbuscular mycorrhizal (AM) fungi form one of the oldest and most widespread obligate mutualisms on Earth, yet they must survive independently while dispersing between hosts. Spores bridge this vulnerable host-free phase, and their morphology should therefore reflect the demands of persistence, dispersal, and establishment. However, the macroevolutionary trajectories of AM spore morphology remain poorly resolved, limiting our ability to determine whether spores diversified into multiple designs or remained constrained around a common architecture. Here, we construct the first quantitative morphospace of AM fungal spores and infer macroevolutionary patterns of trait evolution. We find that AM fungal spores have diversified mainly through scaling rather than redesign. The morphospace is dominated by size, with spore dimensions and wall volume coordinated through near-isometric scaling. Shape remains predominantly near-spherical across sizes, although the largest spores allocate proportionally less material to the wall, while ornamentation and coloration form a largely independent axis of surface variation. Most species occupy a narrow region of trait space, with distantly related lineages converging on similar trait combinations. We propose that adaptive filtering and construction economy jointly maintain this architecture. Near-spherical geometry may provide an efficient solution for packaging and protecting the reserves needed to persist between hosts while minimizing investment in wall material, whereas surface traits may mediate dispersal vectors. Functionally, this architecture suggests that AM fungal spores are shaped more by persistence through time than by dispersal through wind. The AM fungal spore morphospace thus links conserved spore design to the challenge of dispersal in an obligate mutualist.

## INTRODUCTION

Arbuscular mycorrhizal (AM) fungi are obligate symbionts, yet they have retained horizontal transmission throughout their evolutionary history. That is, between hosts, they must persist autonomously as soil-borne spores that survive, disperse, germinate and initiate presymbiotic growth using only the genetic material and reserves packaged before separation from the plant ^1–3^. 400 Ma Fossil evidence indicates that spore production was already present in the earliest known symbiotic associations with plants ^4^. The maintenance of spores among AM fungi from these early associations to all extant AM fungal species shows that these structures provide a successful transmission strategy despite the risks imposed by host absence. This makes the spore stage central to the evolutionary success of the symbiosis. Yet the phenotypic adaptations that make this host-free phase possible remain poorly understood. Because spores encounter diverse abiotic and biotic challenges in soil, their morphology should reflect the demands of protection, persistence, dispersal and establishment. We lack a comparative framework for determining which spore phenotypes have evolved, which trait combinations are constrained, and how these traits have enabled the long-term maintenance of horizontal transmission.

Research on AM fungal phenotypic traits is heavily biased toward the symbiotic phase ^5,6^. The prevalent paradigm is that only traits directly involved in host interaction are considered functional ^7^. This view has been challenged in recent global analyses that show that some spore trait combinations are more prevalent in certain climates than others ^8^. Mapping the phenotypic diversity of AM fungal spore phenotypes provides an opportunity to understand the challenges of dispersal for this mutualism, as spore traits determine dispersal success by determining how resources are packaged, protected and deployed to survive this phase.

Up until recently, the use of spore traits to determine how they survive this phase was limited due to limited amount of data. However, the recent development of the TraitAM database ^9^ overcomes this limitation and enables, for the first time, the construction of a global morphospace of AM fungal spores. These spores comprise two functionally distinct components: a cytoplasmic interior containing the genetic material and reserves needed to initiate presymbiotic growth ^10–12^, and a cell wall that protects these contents and mediates interactions with the external environment ^13^. TraitAM captures variation in both components. Spore size and shape are proxies for the total amount of internal material packaged and its geometry, whereas wall thickness, pigmentation and surface ornamentation describe investment in protection and modification of the spore–environment interface that interacts with the dispersal vector.

Here, we leverage spore trait data to determine the spore construction hypervolume across all extant AM taxa to infer the macroevolutionary forces that drove this variation. Trait hypervolumes represent multivariate projections of all traits within a given phenotype and clade ^14^. This approach allows not only to visualize the full range of trait combinations that are observed in extant taxa ^15^, but also to identify combinations that are absent—revealing the trajectories in the evolution of phenotype, usually in combination with models, capturing a specific evolutionary mechanism. The concept has deep roots in evolutionary biology, dating back to studies linking beak shape and size in Darwin’s finches to adaptation of food preferences ^16^. It has since been widely applied in birds, with recent examples linking egg-trait hypervolumes to flight capacity and morphology-trait hypervolumes to dietary strategies ^17^. This approach is not exclusive to animals; in plants, trait hypervolume analyses have linked flower traits to pollination syndromes ^18^ and leaf traits to axes of resource acquisition and conservation ^19^. Further exploring how traits are combined in these hypervolumes reveal the trait trade-offs and synergies that have shaped the evolution of clade as well as provide mechanistic understanding in functional traits and their influence on community composition and dynamics ^20–22^. In contrast to plants and animals, trait hypervolume approaches have rarely been applied to microorganisms or symbiotic systems, owing both to limited integration across disciplines and to the scarcity of standardized trait data.

By applying hypervolume analysis to spore traits, we quantified variation in spore construction across extant AM fungal species and used this framework to infer the evolutionary processes shaping spore morphology. Specifically, we addressed two interrelated questions. First, has an single spore morphology evolved, or are there multiple phenotypes, and which phenotypic characteristics define these optima ^23^?. Second, which trait synergies and trade-offs structure variation in spore morphology? Overall, our work provides a more holistic understanding of the evolution of complex mutualistic organisms by explicitly considering trait variation across life stages both with and without the host.

## Results

### AM fungi occupy a strongly structured spore morphospace

Across the Glomeromycota phylum containing all AM fungi, variation in spore morphospace was primarily structured by the tight correlation between spore size and cell wall investment (Fig. 1). This main axis of variation was defined by strong positive correlations of the linear dimension of spore volume and wall investment (i.e. spore width, length, and wall thickness, Fig. S1). The distributions of these traits were unimodal in log space and closely approximated a log-normal distribution. The second axis, orthogonal to size dimensions, corresponds to variation in ornamentation and color, indicating that the presence and magnitude of ornamentation and pigmentation is decoupled from variation in spore size. Ornamentation height also exhibited a unimodal distribution (in raw scale), but with pronounced kurtosis, indicating that variation in this trait is restricted to a small group of AM taxa. Spore color, while in ordinal scale, showed that most AM fungi produce light-pigmented spores, with an almost equal split between hyaline and dark-pigmented taxa around light-pigmented ones. These distributional patterns and trait dependencies were supported by hypervolume analyses comparing the observed morphospace against four null models of trait independence and uniform or normal distributions of trait space ^19^. The observed hypervolume differed significantly from all null expectations (p < 0.01, Fig. S2. Table. S1).

**Figure 1.**
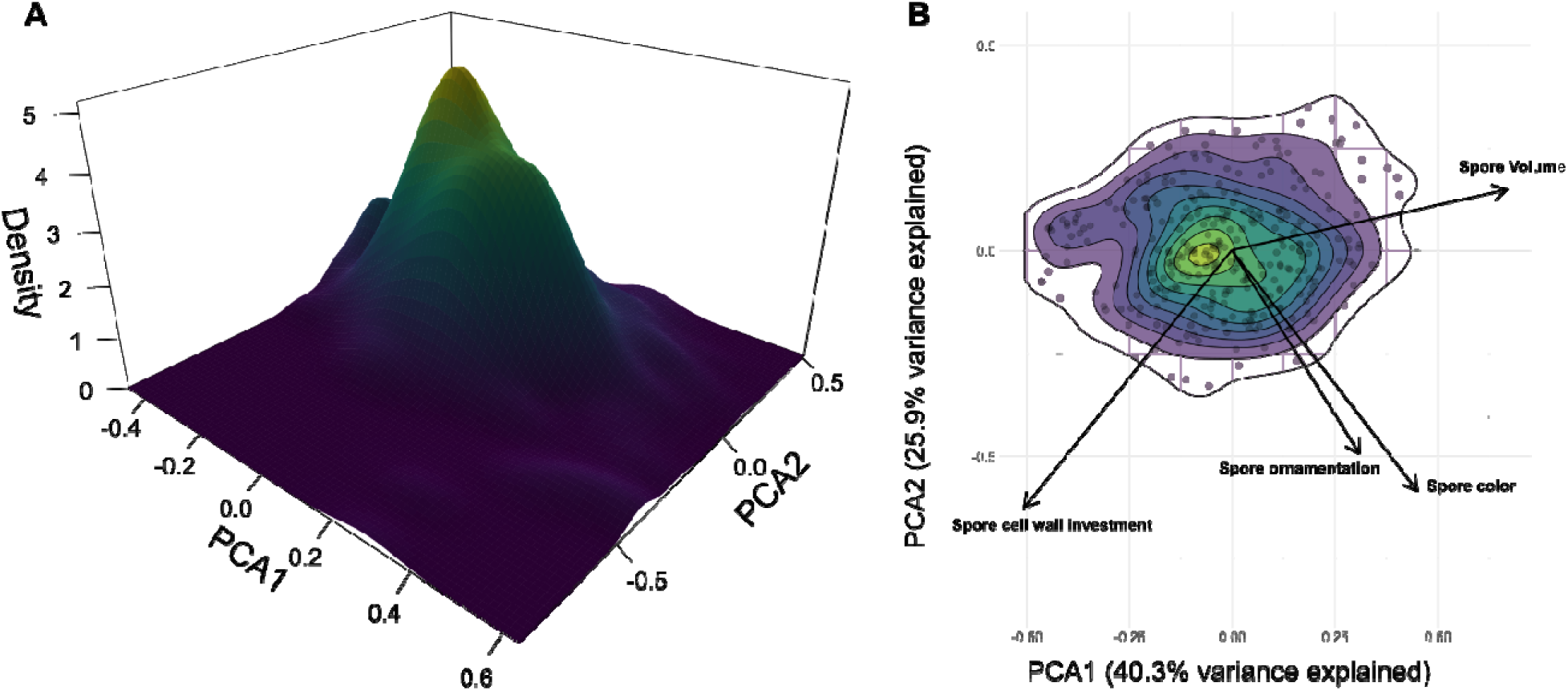
AM fungal spore morphospace. The morphospace is visualized with principal component analysis (PCA) summarizing variation in morphospace based on total volume, investment into cell wall, ornamentation height and color. A) Density plot projection showing the concentration of spore phenotypes around one peak. B) Bi-plot showing the correlation structures of the variable behind the AM fungal spore morphospace.

### AM fungal spores diversified in scale while retaining a common geometry

To infer the mechanisms underlying the tight correlation among spore equatorial and polar axis (i.e. width and length dimensions respectively) and cell wall thickness, we determine the scaling relationship between these traits with total size. We found a nearly isometric relationship in spore dimensions: width and length are tightly correlated at an approximately 1:1 ratio across species, with a constant elongation across total size variation (Fig. 2a). This pattern indicates that, on average, perfect spherical shapes are maintained regardless of the large variation in size with just a handful of species deviating strongly from this trend (Fig. 2a).

**Figure 2.**
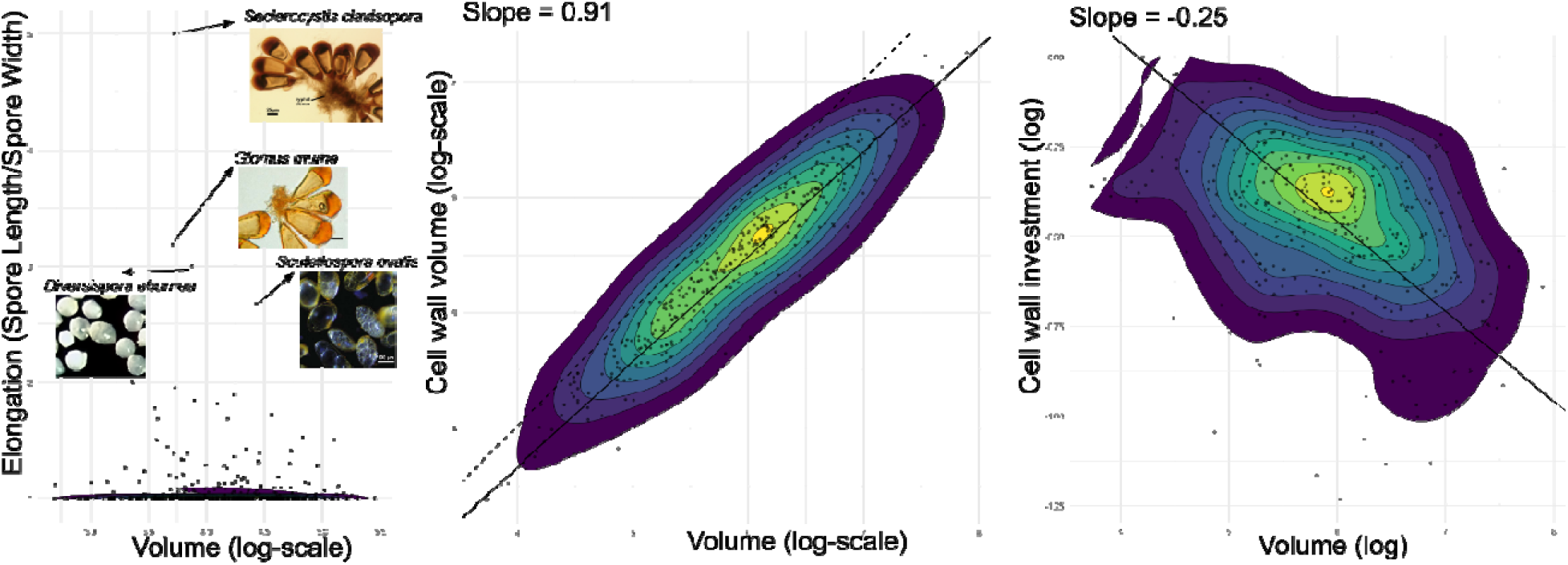
Size–trait relationships closely follow isometric scaling. (A) Across the observed size range, most spores maintain shapes closely approximating perfect spheroids. A small subset of species deviates markedly from this trend; depicted here are the four species that produce spores with the most ellipsoidal shapes (i.e. where spore polar axis is more than two times spore equatorial axis). (B) Cell wall volume increases approximately proportionally with spore size, closely following isometric scaling. However, the large spores show a slight negative deviation from proportionality, exhibiting lower cell wall volumes than expected under strict isometry. (C) Expressed as the ratio of cell wall volume to total spore volume, this deviation becomes more apparent: larger spores allocate proportionally less volume to the cell wall than smaller spores.

We also found a nearly isometric relationship in the volume allocation of cell wall relative to total size. That is, cell wall thickness scales almost proportional to spore length, resulting in a nearly 1:1 relative investment in cell walls across most of the spore size variation across species. However, the slope of the relationship is lower than 1 (Fig. 2b), indicating that as spores attain the largest range of size, the relative allocation into spore volumes could not be maintained. This trend is evident in the negative allometric relationship of cell wall investment (as the ratio of cell wall volume) to total spore volume (Fig. 2c). These patterns suggest a limit in the investment of cell wall in species with extremely large total sizes.

### Evolutionary trajectories of the spore morphospace

Brownian motion (BM) consistently received the least support across all three quantitative spore traits, indicating that morphological divergence did not accumulate progressively alongside the evolutionary diversification of AM fungi. Spore volume was best described by an Ornstein– Uhlenbeck (OU) model (AICc = 462.19), with substantially less support for white noise (WN; AICc = 525.96; ΔAICc = 63.77) and BM (AICc = 566.40; ΔAICc = 104.21). Thus, while spore size retains phylogenetic structure, its divergence remains bounded within a restricted region of trait space rather than increasing continuously among lineages. For both cell-wall investment and ornamentation height, OU and WN models received similar support. Cell-wall investment was best described by WN (AICc = −155.75), closely followed by OU (AICc = −153.35; ΔAICc = 2.40), whereas BM received substantially less support (AICc = 149.68; ΔAICc = 305.43). Likewise, ornamentation height was similarly supported by WN (AICc = 821.18) and OU (AICc = 822.62; ΔAICc = 1.44), while BM again performed substantially worse (AICc = 1186.73; ΔAICc = 365.54). Together, these results consistently show that variation in spore morphology does not track the progressive diversification of AM fungal lineages: spore size remains phylogenetically structured but evolutionarily bounded, whereas variation in cell-wall investment and ornamentation shows little evidence of accumulating with shared evolutionary history.

**Figure 3.**
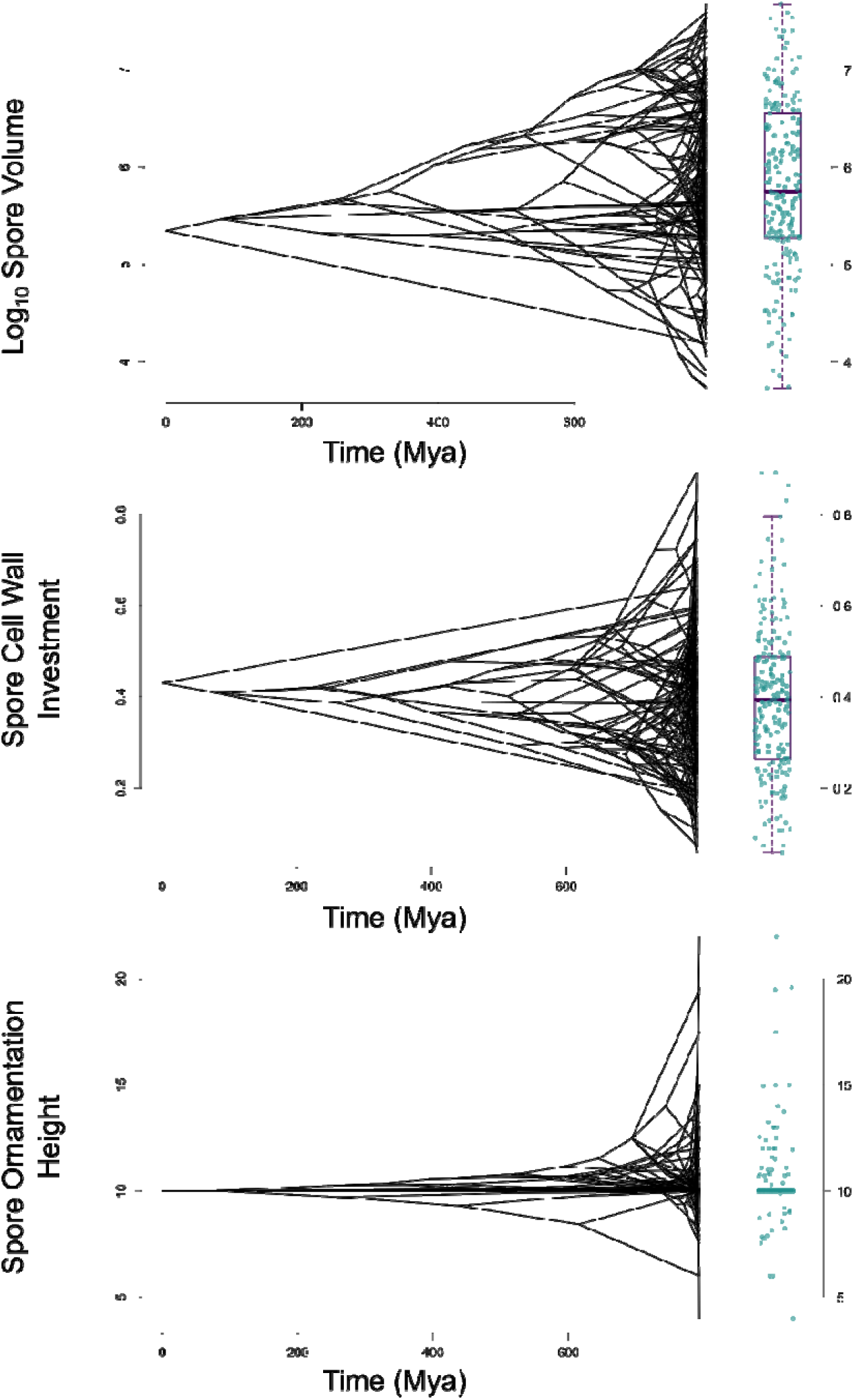
Phenograms showing the evolution of three AMF spore traits across the time-calibrated phylogeny. Each panel maps extant species (tips) and their inferred ancestral values (internal nodes) onto a time axis derived from the penalized likelihood chronogram (based on node estimates as in ^24^). The horizontal axis represents absolute divergence time (millions of years ago, Mya), and the vertical axis represents trait value. The marginal distribution on the right of each panel shows the spread of extant trait values as a boxplot with jittered individual species. **(a)** Log□□-transformed spore volume (µm³). Ancestral lineages show a wide range of volumes that converge toward intermediate values in more recent clades, suggesting neither strong directional drift nor tight constraint. **(b)** Cell wall investment (shell volume / total volume, unitless). Values are tightly clustered across most of the phylogeny, with low variance among both ancestral and extant lineages, suggesting cell wall variation is decoupled from diversification patterns in AM fungi. **(c)** Spore ornamentation height (µm; baseline = 10 µm, corresponding to smooth spores). The majority of lineages are clustered at the baseline, with sporadic independent shifts to higher ornamentation in scattered clades, indicating that ornamentation arises repeatedly but is not broadly conserved across the phylogeny.

## Discussion

The AM fungal spore morphospace reveals strongly constrained evolution of spore morphology, characterized by the retention of a common architecture rather than unrestricted diversification into alternative propagule designs. Unlike morphospaces of macro-organisms such as plants, which often exhibit multiple peaks corresponding to distinct trait combinations ^19^, the AM fungal spore morphospace is predominantly unimodal, while phylogenetic models consistently reject the progressive accumulation of morphological divergence expected under Brownian motion. Thus, the diversification of AM fungal lineages was not accompanied by a corresponding diversification of spore morphology. Instead, despite about four orders of magnitude variation in total size, spore dimensions and wall volume scale in a tightly coordinated manner, while a near-spherical shape is retained as the predominant geometry across the group. This size-dominated axis of diversification therefore reflects changes in scale rather than redesign of the underlying architecture. Ornamentation and coloration introduce a second, largely independent axis of variation, adding surface-level novelty without disrupting this core design.

### Adaptive filtering and construction economy as complementary explanations for conserved spore architecture

Two non-mutually exclusive mechanisms may explain the restricted architecture of AM fungal spores. The first is adaptive filtering. Spores must support dormancy, survival, dispersal, germination, and establishment during the host-free phase, and deviations in their design may therefore carry direct fitness costs. Traits with direct fitness consequences often show limited variation because even small departures from optimal values are less fit ^25^. The concentration of lineages around similar trait combinations is therefore consistent with a common adaptive region in which only a limited set of phenotypes performs reliably. Alternative phenotypes may arise developmentally, as suggested by within-species variation reported in AM spore descriptions ^9^, but may rarely persist over evolutionary time. Rather than promoting broad morphological divergence, the multiple demands placed on spores may therefore funnel evolution towards a comparatively restricted set of successful phenotypes.

Similar patterns of constrained morphological evolution have been documented for structures functionally analogous to AM spores. In bird eggs, for example, shell thickness is tightly linked to fitness because it must remain within a range that provides sufficient protection while still permitting embryonic development and successful hatching ^26,27^. In fact, AM fungal taxa, similar restrictions to cell wall investment may exist above which investment in cell wall may jeopardize successful germination. Ecological transitions can also produce repeatable changes in fungal spore morphology. Ectomycorrhizal fungi produce, on average, larger and more ornamented spores than their saprotrophic relatives^28^ suggesting that the demands associated with symbiosis repeatedly favour particular regions of spore morphospace. Together, these comparisons support the broader idea that propagule morphology is filtered by the conditions under which survival and establishment must occur.

The second mechanism is construction economy. The two principal components of an AM fungal spore (i.e. the reserve-rich interior and the enclosing cell wall) differ in their resource origins and biosynthetic requirements. Much of the lipid stored within the spore ^12^ is supplied by the host plant ^29^, whereas the wall consist of complex fungal biomolecules that must be synthesized by the fungus ^30–33^. Increasing internal volume and increasing wall material should therefore impose different resource demands. This asymmetry provides a possible explanation for both the broad variation in spore size and the persistence of spherical geometry ^34^. For a given wall thickness, a sphere encloses the greatest volume with the least surface area, minimizing the material required to protect a given amount of internal content. Larger spores can consequently package more reserves, whereas departures from sphericity would increase the wall material (and therefore the fungal investment) required to enclose them. Assuming constant biochemistry of spores across all AM taxa (an assumption that remains to be tested), the widespread retention of near-spherical spores is thus consistent with selection favouring geometries that maximise internal volume while limiting wall construction.

The slight negative allometry further supports resource constraints on spore construction, as it indicates that species producing spore in the largest size ranges invest proportionally less in wall material than expected under isometric scaling. This pattern suggests that as spores increase in size—reaching the largest known in the fungal kingdom—the costs of wall construction progressively limit further investment. Even with an allometric slope close to one (0.92), its effect accumulates across the four-order-of-magnitude range in spore volume, reducing the predicted proportion allocated to the wall from approximately 50% in the smallest-spored taxa to 25% in the largest. Because spherical geometry is already the most efficient shape to minimize wall allocation of material for a given volume, deviations of this shape would only incur higher resource costs. As a result, large-spored lineages may approach a constructional boundary beyond which additional wall investment becomes metabolically prohibitive. Direct measurements of spore construction costs are needed to determine whether energetic limitation in responsible for the observed allometry.

Similar allometric patterns occur in other propagules. In plant seeds, coat thickness—the functional analogue of the AM fungal spore wall—scales negatively with seed size across wild species, crops and plant families ^35^ ^36^ ^37^. Thus, as in AM fungi, large-seeded species therefore allocate proportionally less material to protective structures than smaller-seeded species. Likewise, butterfly species producing larger eggs allocate proportionally less nitrogen per offspring than species producing smaller eggs ^38^. These comparisons suggest a broader limitation in the construction of propagules as they increase in size: investment in protection layers increase more slowly than total propagule volume and possibly an indication of a limit in the effectiveness of protection.

We emphasize that both of these mechanisms, adaptive filtering and construction economy, act as complementary explanations. That is, the same architecture may be favored because it is both functionally optimal and economical to construct. Thus, stabilizing selection and resource constraints may act in the same direction, repeatedly drawing lineages towards a common region of morphospace. Distinguishing their relative contributions will require more evidence to extablish the links between spore morphology, construction costs, survival, germination and establishment success across a suite of conditions.

### AM fungal spore morphospace more consistent dispersal through time rather than by wind

The AM fungal spore morphospace suggests is more congruent with a structure that serves for dispersal through time than dispersal by wind. In many wind-dispersed spores, their morphology is shaped by the need to break the still-air boundary layer surrounding them reach dispersive airflows. As a result several mechanism to forcibly ejected spores have arisen, alongside repeatedly evolved drag-minimizing spheroidal shapes (bullet-like) that increase their initial flight distance ^39,40^. AM fungal spores exhibit a contrasting suite of traits: they are exceptionally large, predominantly near spherical that do not minimize drag, and a lack of a specialized launch mechanism. Although AM fungal spores have recovered from air, particular small-spored taxa ^41^, wind is unlikely the primary force shaping the evolution of the spore architecture. Instead, we hypothesize that this architecture is shaped mainly by dispersal through time—the capacity to remain viable between one host and the next. That capacity of spores to endure without a host is remarkable, for instance, AM fungal spores have been found viable even after 6 years without the host in artic environments ^42^. Large size provides the reserves required for survival, germination and presymbiotic growth, while spherical geometry encloses and protects those reserves economically. Variation in size may therefore represent different levels of investment in persistence and establishment rather than alternative solutions for aerodynamic transport.

Cell wall traits, in contrast, may carry a stronger signature of dispersal through space with specialized vectors. Ornamentation could influence adhesion to soil particles or animal vectors, whereas coloration may indicate differences in wall chemistry that affect resistance to desiccation, microbial attack or passage through digestive systems. Animal-mediated dispersal provides a plausible selective context, as viable AM fungal spores can survive passage through mammalian digestive tracts ^43,44^). Under this view, the conserved core architecture supports persistence through the host-free dispersal, whereas variation at the spore surface may mediate movement and survival across particular dispersal adaptations. The uneven distribution of these traits—and particularly the restriction of pronounced ornamentation to a small subset of taxa— suggests that surface differentiation represents lineage-specific departures layered onto an otherwise conserved design.

Thus, while AM fungal spore evolution may be constrained by an architectural core design, surface modifications that provide greater scope for ecological specialization. For example, larger spores are more prevalent in warm, wet climates but are associated with smaller geographic range sizes, suggesting a trade-off between investment in persistence and broad dispersal potential ^8^. Similarly, cell wall investment increases toward cooler, drier climates and is the strongest morphological predictor of species range size, with intermediate levels of investment associated with the broadest geographic distributions. Thus, evolutionary constraint on the overall architecture does not imply ecological equivalence among spore phenotypes but instead, the observed variation within the morphospace may contribute to how AM fungi persist and disperse across contrasting environments.

### Outliers to the AM geometric design are a window into distinctive ecologies

Several species depart substantially from the dominant scaling relationships and morphospace regions identified here. These outliers show that alternative morphologies are evolutionarily attainable, even if rarely realized. Understanding why some AM taxa produce strongly ellipsoidal spores or allocates more wall volume than expected for their size could therefore reveal ecological pressures or life-history strategies ^45^ that differ from those shaping most of the group. Comparable deviations from general allometries have exposed specialized functions elsewhere: the disproportionately large bill of toucans relative to allometric relationship of bird size, for example, is explained by its function to dissipate heat ^46^. For AM taxa, information about the chemistry of the wall of those spores would provide valuable information as it would tell whether the species that invest in large spores, or they produce spheroidal shapes that require higher allocation in cell wall can do it because they use metabolically less expensive biochemistry. Likewise, information on their habitat and natural history would be essential to disentangle the ecological and evolutionary circumstances associated with these exceptional phenotypes represent an important next step towards understanding the processes governing morphospace occupation.

### The spore morphospace links with the maintenance of AM symbioses

When placed in a broader evolutionary context, even the smallest AM fungal spores in this morphospace are approximately two orders of magnitude larger than most other fungal spores ^47^. Thus, we hypothesize that the maintenance of consistently large sizes represents an evolutionary lower value imposed by horizontal transmission ^3^. As obligate symbionts, AM fungi must package sufficient reserves and nuclei within each spore to remain viable, germinate and initiate presymbiotic growth during the uncertain interval between hosts ^10^. This requirement may impose a minimum internal volume below which successful transmission becomes unlikely. At the same time, these reserves and the genetic material they support must remain protected from the external environment until a suitable host is reached. The predominance of spherical geometry may provide an economical solution to this dual requirement, maximizing the internal volume available for reserves while minimizing the wall material needed to enclose and protect them. Thus, AM fungal spores may diversify above this minimum size but remain constrained to a common geometry that balances storage with protection. Ornamentation and coloration may then provide a more flexible surface through which spores adapt to transmission environments without altering this fundamental design. The climatic and geographic associations of these same traits reinforce this interpretation that variation within the conserved spore architecture is associated with both environmental conditions and species geographic distributions ^8^.

The evolutionary persistence of the AM symbiosis depends on resolving a fundamental paradox: fungi that rely entirely on their plant hosts for their basic metabolism must nevertheless endure dispersal phase decoupled from host. Spores bridge these two phases, carrying the resources to start a new symbiotic phase through an interval of temporary independence. Our results suggest that this transition has not resulted in the evolution of many radically different propagule designs, but through scaled-dependent variation that maintains a remarkably conserved architecture. Differences in size and surface properties may represent alternative transmission strategies, yet these remain bounded by a common geometry and economy of construction. Whether these boundaries reflect the energetic demands of building a viable spore, selection for reliable survival and establishment, or an interaction between both remains to be determined. Viewed in this way, the AM fungal spore morphospace is not simply a description of morphological diversity; it is an evolutionary record of the solutions that have allowed an obligate symbiosis to persist across hosts, environments, and more than 400 million years of change.

## Material and methods

### Trait data and spore morphology calculations

Spore morphological data were sourced from the TraitAM database ^9^. For each species, two linear dimensions (polar axis and equatorial axis) were recorded as ranges (minimum, mean, and maximum), along with cell wall thickness (minimum, mean, and maximum) and descriptors for spore color and ornamentation height. The polar axis was defined as the longest axis of the spore; when both axes were equal the spore was treated as a perfect sphere. This convention restricts spore shape to either spherical or prolate spheroid forms, excluding oblate (saucer-shaped) morphologies.

Spore volume was calculated using the formula for a prolate spheroid:

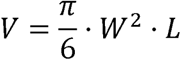

where *W* is the equatorial (width) axis and *L* is the polar (length) axis. The inner volume—the space enclosed by the cell wall—was calculated by subtracting twice the mean wall thickness from each axis before applying the same formula. Cell wall volume (shell volume) was then the difference between total volume and inner volume, and cell wall investment was defined as the ratio of shell volume to total volume.

Spore ornamentation was measured as mean ornament height (*μ*m). Because the majority of AMF species produce smooth spores (ornamentation height = 0), a constant of 10 *μ*m was added to all values prior to analysis. This rescaling shifts smooth spores to a baseline of 10 and preserves the relative differences among ornamented species while avoiding zero-inflation issues in log-space analyses. Species lacking measurements for wall thickness or spore dimensions were excluded from downstream analyses; four additional species with missing ornamentation data were excluded only from ornamentation-specific analyses.

### Measurement of AM fungal spore morphospace

To characterize the multivariate trait space (i.e. the morphospace) occupied by AM fungal spores, a principal component analysis (PCA) was performed on four quantitative traits: log_10_-transformed spore volume, log_10_-transformed cell wall investment, spore color (ordinal ranking ranked 0–6 from hyaline to heavily melanized), and mean ornamentation height. The PCA was computed using the rda() function from **vegan**^48^ with standardization (correlation matrix), so that all traits contributed equally regardless of their original scale. Spore shape was not included as it remained mostly constant across all species and thus, did not contribute to total variation in the morphospace. The observed morphospace was visualized using bi-plots overlaid with density projections.

### Estimations of deviations of observed morphospace to null-models of multi-trait evolution

To evaluate whether the observed AM fungal spore morphospace is more constrained than expected by chance, the volume of the observed trait space was compared against four null models of increasing biological realism, following the framework of ^19^. Trait space volume was quantified as the volume of the convex hull enclosing the central 95% of species in the standardized three-dimensional PCA space (PC1–PC3), computed using convhulln() from the **geometry** package. The 95% subset was defined by retaining species whose squared Euclidean distance from the centroid fell below the 95th percentile, implemented via the subselect.data() utility function. PCA was performed using dudi.pca() from **ade4**^49^.

For each null model, 999 permuted datasets were generated, their convex hull volumes calculated, and the observed volume was tested against the resulting null distribution using a one-sided randomization test (as.randtest(), **ade4**), with the alternative hypothesis that the observed volume is smaller than expected by chance (i.e., the morphospace is more constrained). The four null models were:

Null Model 1. Uniform distribution, orthogonal traits. Each trait was independently resampled from a uniform distribution spanning the observed minimum to maximum of that trait. This is the most conservative null: it assumes no preferred trait values and no correlations among traits, producing a hypercube-shaped space that is the maximum possible volume given the observed ranges. Null Model 2. Normal distribution, orthogonal traits. Each trait was independently drawn from a normal distribution with mean zero and the observed standard deviation of that trait. This model allows for a concentration of species near mean trait values but still assumes all traits evolve independently with no inter-trait correlations. Null Model 3. Observed marginal distributions, orthogonal traits. This model preserves the exact empirical marginal distribution of each trait—including any skewness or multimodality—while eliminating all inter-trait correlations. It therefore tests whether the observed volume is smaller than expected if traits were free to vary independently within their real ranges. Null Model 4. Normal distribution, observed correlation among traits. Independent standard normal variates were drawn for each trait but maintaining the same pairwise correlations as the observed. Thus, this model assume a process of where the evolution of each trait is not independent from each other.

For each null model, the percentage reduction in morphospace volume relative to the null expectation was also computed as 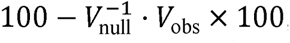, providing an intuitive measure of how much smaller the realized trait space is compared to each null scenario.

In addition to convex hull volume, the concentration of species within the trait space was assessed by comparing observed versus null cumulative species-accumulation curves across equal-sized hypercells in PCA space, visualized for all four null models.

### Allometric scaling of spore dimensions

To evaluate whether spore dimensions scale isometrically or allometrically across species, pairwise scaling relationships among spore traits were tested using Standardized Major Axis (SMA) regression implemented in the **smatr** package ^50^. SMA regression is preferred over ordinary least squares for allometric analyses because it treats both axes as measured with error and seeks the line of best fit through the major axis of the joint distribution, rather than minimizing residuals in only one direction^51^. All regressions were performed on log_10_-transformed values.

Three focal relationships were examined and visualized. First, elongation (polar/equatorial ratio) was regressed against spore size (polar × equatorial area), with a null slope of zero tested (isometry would imply shape is independent of size). Second, shell volume was regressed against total volume on log–log axes, testing a null slope of 1 (strict isometry). Third, the ratio of shell volume to total volume (investment) was regressed against total volume on log–log axes, testing a null slope of 0 (investment independent of size).

### Phylogenetic analyses

The phylogenetic tree used in all comparative analyses was the TraitAM consensus phylogeny, read in NEXUS format using read.nexus() from **ape** ^52^. Tip labels were standardized to match the taxonomy in the trait dataset using a series of regular expression substitutions, and two outgroup taxa (*Arabidopsis thaliana* and *Oryza sativa*) were removed with drop.tip().

The topology-only tree was converted to an ultrametric, time-calibrated chronogram using the penalized likelihood method implemented in chronos() from **ape**, with a correlated-rates clock model (model = “correlated”). This model assumes that evolutionary rates are autocorrelated between parent and daughter branches, which is appropriate for a group like AMF where rate variation is expected to accumulate gradually. We time-calibrated the phylogeny using two node-age constraints: the split between Dikarya and non-Dikarya lineages (796–1253 Ma) and the divergence between Gigasporales and Glomerales (408–580 Ma). These bounds were derived from published molecular-clock estimates ^53^ and fossil-calibrated fungal phylogeny ^24^. Polytomies introduced during the standardization step were resolved randomly using multi2di(), and zero-length branches were assigned a minimal positive length (10□□) to avoid numerical failures in downstream likelihood calculations.

Evolutionary patterns of continuous trait variation were visualized as phenograms (also known as traitgrams) using phenogram() from **phytools** ^54^. A phenogram projects species onto a phylogeny such that the horizontal axis represents geological time and the vertical axis represents trait value, allowing visual assessment of whether trait disparity accumulates gradually (Brownian Motion pattern) or is constrained toward a central value (Ornstein-Uhlenbeck pattern). Phenograms were produced for three traits: log spore volume, cell wall investment, and spore ornamentation height. Each phenogram was paired with a marginal boxplot and jitter strip showing the distribution of extant trait values, and exported as a PDF using a Cairo device to ensure font embedding (important for figure editing in vector graphics software).

To formally evaluate the mode of trait evolution for each continuous trait, three evolutionary models were fitted to the time-calibrated tree using fitContinuous() from **geiger**^55^:

- **Brownian Motion (BM)**: trait values evolve as a random walk, with variance accumulating linearly with time. This is the null model of neutral drift.
- **Ornstein-Uhlenbeck (OU)**: trait values are subject to a deterministic pull toward an optimum (*θ*), with the strength of this pull controlled by the parameter *α*. Higher indicates *α* stronger stabilizing selection.
- **White Noise (WN)**: trait values are drawn independently from a distribution with no phylogenetic structure, serving as a null for the complete absence of phylogenetic signal.

Model fits were compared using AIC (Akaike Information Criterion) values returned by fitContinuous(). Additionally, phylogenetic signal was quantified for each trait as Pagel’s *λ* using phylosig() from **phytools**, with a likelihood ratio test against the null hypothesis *λ* = 0 (no phylogenetic signal). *λ* ranges from 0 (trait evolution independent of phylogeny) to 1 (trait evolution fully consistent with Brownian Motion expectations). Species with missing trait data were excluded from the relevant model, and the tree was pruned accordingly by treedata() from **geiger** prior to model fitting.

All analyses were conducted in R (R Core Team) and **ggplot2** for all figures.

## DATA AVAILABILITY

TraitAM, a global spore trait database for arbuscular mycorrhizal fungi, is deposited in the Dryad Digital Repository at https://doi.org/10.5061/dryad.6hdr7sr8z. It is publicly available for download under the CC0 public domain dedication given proper scholarly citation of the version used.

## CODE AVAILABILITY

Code for calculation of spore trait metrics and LSU-based Glomeromycotinan phylogeny construction is deposited in the Dryad Digital Repository at https://doi.org/10.5061/dryad.6hdr7sr8z. Code and data needed for reproducing the all analysis and Figures in this manuscript is found in the Github repo: https://github.com/aguilart/AM-fungal-morphospace

## AUTHOR CONTRIBUTIONS

VBC and CAA-T conceived the study. VBC, CAA-T, MCR and JB prepared the initial manuscript. CAA-T, SPL and VBC developed subsequent versions. CAA-T conducted statistical and phylogenetic analyses, with LN performing the hypervolume analyses. All authors contributed to revising the final manuscript.

## ACKNOWLEDGEMENTS

This work was supported by the National Science Foundation (DEB-2205650) and Dartmouth College. CAA-T acknowledges funding by the Research Council of Finland (Decision number 356191).

## Notes

### Competing Interest Statement

The authors have declared no competing interest.

