## Supplementary material for "The first spectrum of spore form and function reveals constrained evolution in mycorrhizal symbiosis": Supplemntary Figures and Table

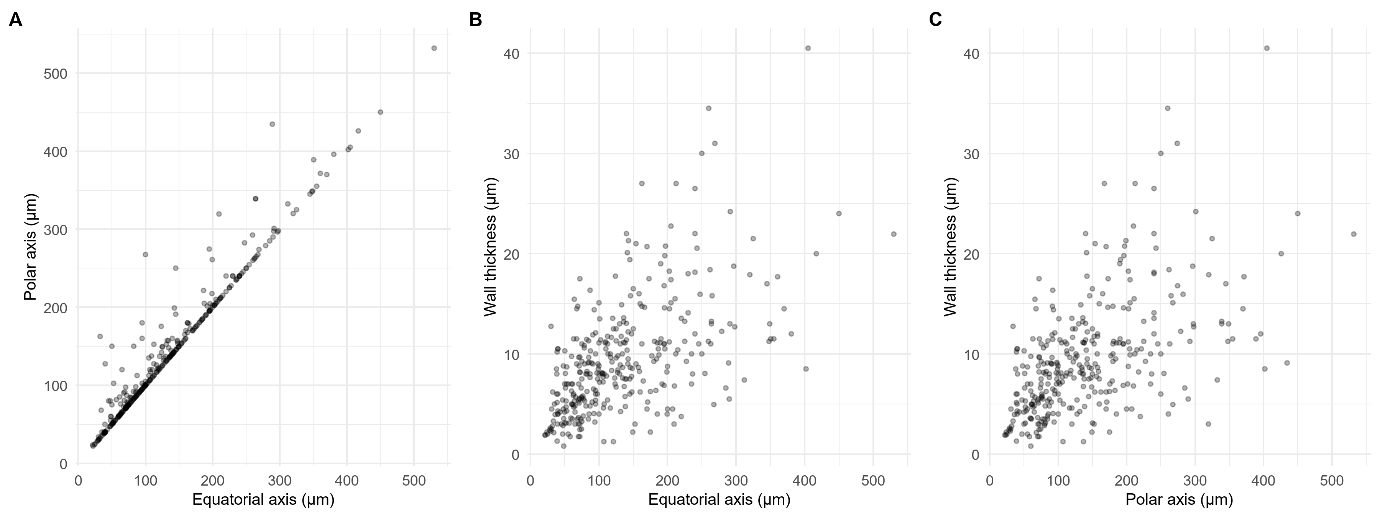

**Figure S1. Pairwise relationships among the three linear dimensions used to calculate spore volume and cell wall investment.** Each panel shows species mean values (points) overlaid with filled 2D kernel density contours (shading proportional to local species density). **(A)** Equatorial (spore width) axis vs. polar (spore length) axis. Because the polar axis is constrained to be ≥ the equatorial axis to reflect prolate spheroid shape of AM spores, all species fall on or above the 1:1 line, with the majority clustered near spherical morphologies. **(B)** Equatorial axis vs. mean cell wall thickness. **(C)** Polar axis vs. mean cell wall thickness.

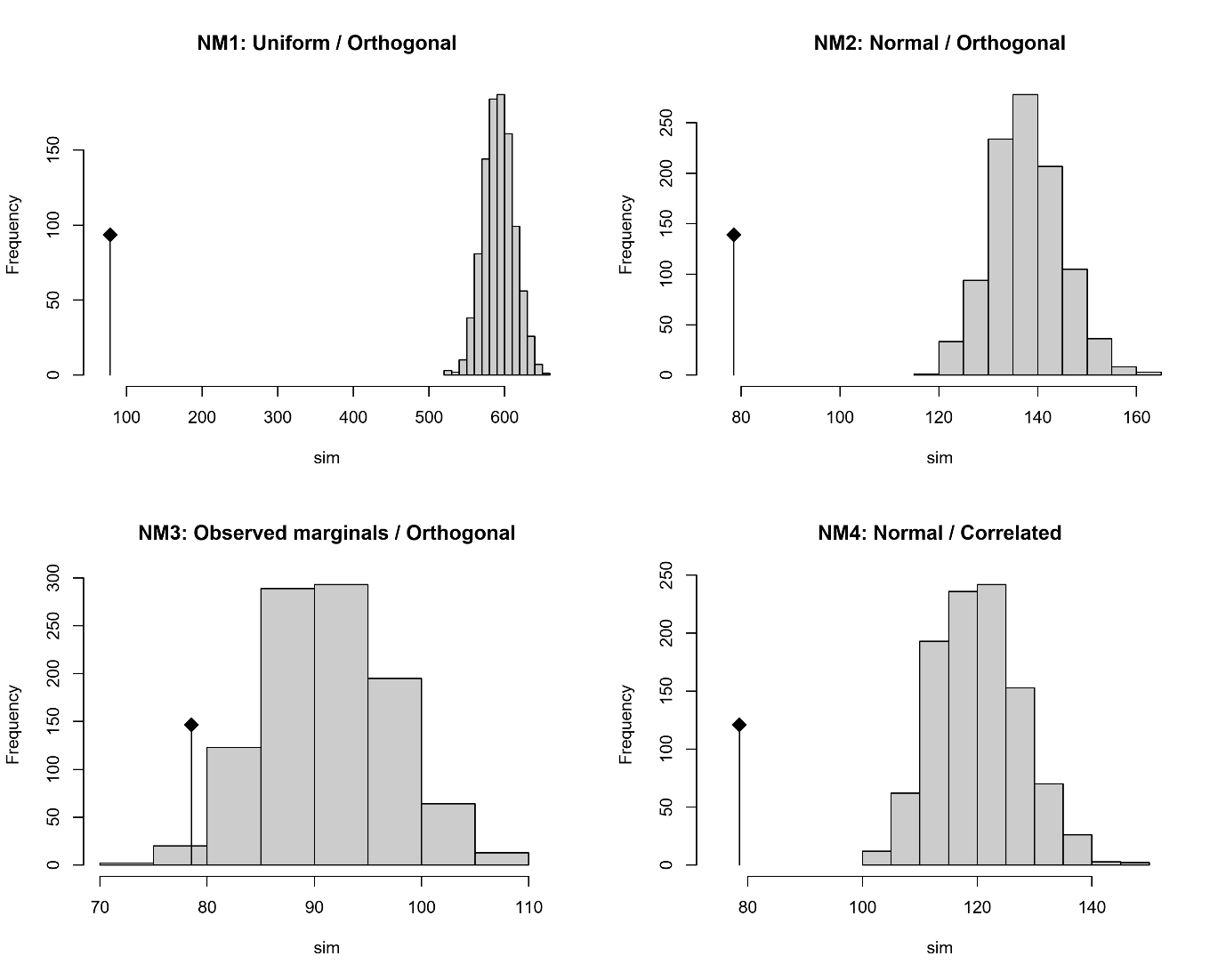

**Observed AMF spore morphospace volume compared to four null model expectations.** Each panel shows the null distribution of 3D convex hull volumes (grey histogram, 999 permutations) alongside the observed volume (red vertical line). Hull volume was computed on the central 95% of species in standardized trait space (log₁₀ spore volume, log₁₀ cell wall investment, spore color, ornamentation height). The alternative hypothesis is that the observed volume is smaller than expected by chance (one-sided randomization test). **Null model 1 (NM1)**: traits drawn independently from a uniform distribution spanning the observed range (hypercube null). **Null model 2 (NM2)**: traits drawn independently from a normal distribution. **Null model 3 (NM3)**: observed marginal distributions preserved but with no inter-trait correlations structure. **Null model 4 (NM4)**: traits drawn from normal distributions with the observed pairwise correlation structure. *p*-values from randomization tests are reported in Table S1.

**Table S1. Comparison of observed AMF spore morphospace volume to four null model expectations.** Observed convex hull volume encloses the central 95% of species in the four-dimensional standardized trait space (log₁₀ spore volume, log₁₀ cell wall investment, spore color, ornamentation height). For each null model, 999 permuted datasets were generated and their convex hull volumes calculated. Null Mean Volume and Null SD Volume summarize the resulting null distribution. Reduction (%) is the percentage by which the observed morphospace is smaller than the null mean, calculated as $100-\bar{V}_{\text{null}}^{-1}\cdot V_{\text{obs}}\times100$. *p*-values come from one-sided randomization tests with the alternative hypothesis that the observed volume is smaller than expected by chance. NM1: traits resampled from a uniform distribution and no trait structure (orthogonal traits); NM2: traits drawn from normal distribution and no trati structure (orthogonal traits); NM3: observed distributions are preserved but no trait structure (orthogonal traits); NM4: traits drawn from normal distribution with observed trait correlation structure.

| **Null_Model** | **Observed Volume** | **Null Mean Volume** | **Null SD Volume** | **Reduction (%)** | **p -value** |
| --- | --- | --- | --- | --- | --- |
| NM1: Uniform / Orthogonal | 78,5261 | 592,1755 | 20,2013 | 86,7 | 0,001 |
| NM2: Normal / Orthogonal | 78,5261 | 137,7126 | 7,1302 | 42,8 | 0,001 |
| NM3: Observed marginals / Orthogonal | 78,5261 | 91,3234 | 5,9961 | 13,6 | 0,011 |
| NM4: Normal / Correlated | 78,5261 | 120,0899 | 7,4739 | 34,4 | 0,001 |
